# Microdroplet-enabled high-throughput cultivation of vaginal bacteria using cervicovaginal fluids

**DOI:** 10.1101/2023.09.26.559375

**Authors:** Corine M. Jackman, James Y. Tan, Xiaoxia Nina Lin

## Abstract

The human vaginal microbiome (HVM) is closely associated with the health of the host. In particular, bacterial vaginosis, a condition where vaginal lactobacilli are reduced dramatically by an overgrowth of various other bacteria, has been linked to increased risk of sexually transmitted infectious diseases, including HIV, and preterm birth. Recent culture-independent studies leveraging next-generation sequencing technology have revealed that the HVM composition differs between women and changes over time. However, questions remain as to the underlying mechanisms and culture-dependent studies are needed for further elucidation of the HVM’s genotype-phenotype relationships and system-level properties *in vivo*. In this work, we have adapted a previously developed microdroplet-based high-throughput cultivation platform for the investigation of vaginal bacteria using the cervicovaginal fluid (CVF) as cultivation medium. Using undiluted CVF collected with Softdiscs discs, we observed the growth of *L. iners* in microdroplets containing CVF pooled from samples with a high prevalence of *L. crispatus*. Although demonstrated with *L. iners*, this work establishes a new framework for culturing microorganisms under clinically-relevant conditions *ex vivo* using minute volumes of host fluids; it can be further extended and adapted for addressing numerous questions about the HVM and other complex microbiomes.

## Introduction

The human vaginal microbiome (HVM) is an essential factor impacting the health of women. Recent culture-independent studies leveraging next-generation sequencing technology have revealed that the HVM composition differs between women and changes over time. It is not clear what factors cause these differences and drive the HVM’s temporal changes. Dysbiosis of the HVM is a hallmark of bacterial vaginosis (BV), a condition where lactic acid bacteria, particularly Lactobacillus spp., is reduced dramatically by an overgrowth of various other bacteria. As the most common lower genital tract disorder of reproductive-age women, BV has been linked to increased risk of sexually transmitted infectious diseases, including HIV, and preterm birth. The HVM is highly complex, harboring a large number of bacterial species. Vaginal lactobacilli make up four of five community types in the HVM. The protective lactobacilli include *L. jensenii, L. gasseri*, and *L. crispatus*, whereas the role of *L. iners* has been unclear.^1^ The fifth community type, composed of diverse strict anaerobes, is associated with BV.^2^ The mechanism behind the shifting of vaginal lactobacilli and the onset of BV still remain largely unknown. One hypothesis for the former is conditional differentiation, which consists of species partitioning their shared niche spaces to survive.^3^

Before and in parallel to culture-independent studies propelled by the advance of sequencing and other technologies in recent years, researchers have also been investigating the HVM using culture-based methods. One critical factor in these studies is the formulation of cultivation media. A major effort has been simulating the cervicovaginal fluid (CVF) experienced by the HVM. CVF is a complex mixture of high and low molecular weight compounds resulted from plasma transudation that filters through the vaginal wall, secretions of Bartholins’ Skeene’s glands, cervical mucus, endometrial and tubal fluids, and other components. CVF is composed of 90% water, organic and inorganic salts, urea, monosaccharides and polysaccharides, mucins, short chain fatty acids, proteins, immunoglobulins, and other macro-molecules. Although many molecules in CVF have been identified there are still some that have yet to be discovered.^4^ Over the years, a number of chemically defined medium have been developed that simulates different facets of vaginal secretions.^5–8^ Dorr *et al*. designed an artificial vaginal fluid (AVF) that mimics the composition of electrolytes, nitrogenous substances, and pH conditions for the purpose of simulating the intravaginal release of retinoids.^8^ Geshnizgani and Onderdonk developed a chemically defined medium (CDM) to grow vaginal isolates by adjusting the pH and final concentrations of nutrients from vaginal secretions, and by supplementing additional components to support growth.^6^ Owen and Katz designed a vaginal simulating fluid (VSF) to imitate pH and osmolarity, which are properties that govern interactions between vaginal fluid and topical contraceptives, prophylactics, and therapeutic products.^7^ Tomás and Nader-Macías also designed a medium simulating vaginal fluid (MSVF) to examine the behavior of six potentially probiotic vaginal lactobacilli at pH 4.2 while supplementing additional nutrients.^5^ Each of these broth medium serve different purposes, have different formulas, but lack all components that have been detected in vaginal fluid. In addition, none of them have been shown to support the growth of the five community state types, and half of them were published before the discovery of *L. iners*, a controversial and frequently detected bacterium in the human vagina.

In 1999, Falsen *et al*. discovered an isolate of *L. iners* after streaking human urine onto blood agar and incubating under anaerobic conditions.^9^ The word “iners” translates from Latin to inactive or inert, which may reflect its nutrient exigent nature and its small genome that is similar to a symbiont or a parasite.^9,10^ *L. iners* does not grow in De Man, Rogosa and Sharpe (MRS) media like other lactobacilli, but has been reported to grow planktonically in New York City (NYC) III broth, and MRS-NYC III (MNC), which both contain horse or fetal bovine serum, respectively.^9,11^ Although MNC grows the four lactobacilli, its formula includes amino acids, peptides, carbohydrates, vitamins, and other essential factors from sources that are foreign to the vagina such as porcine pancreas, bovine skeletal muscle and yeast cells.

Recognizing the deviation of laboratory cultivation media from the *in vivo* environment of the HVM, researchers have also attempted to utilize CVF in experimental studies. For instance, Valore *et al*. examined the antimicrobial activity of vaginal fluids, against resident and non-resident bacteria.^12^ This study reported that five of five donors promoted the growth of *L. crispatus, L. vaginalis*, and *C. albicans*, while *Streptococcus, E. coli*, and *L. jensenii* grew in vaginal fluids from three of five donors. Differences in bacterial fitness allude to heterogeneity in the composition of vaginal fluids across donors. In addition, concentrated vaginal fluids had higher antimicrobial activity against *E. coli*.^12^ Others studies have conducted proteomic analysis on CVF to identify biomarkers for antimicrobial activity, reproduction, forensics, and pathologies.^13^

In this work, we have adapted a previously developed microdroplet-based high-throughput cultivation platform for the investigation of vaginal bacteria using CVF as cultivation medium. Microfluidically generated droplets are water-in-oil emulsions of typically pico- to nano-liter volumes that have been utilized to cultivate and characterize an increasing number of microbial systems.^14–18^ Cultivating vaginal bacteria in microdroplets that contain CVF offers the possibility of examining the physiology of vaginal bacteria in small volumes of host fluid mimicking the *in vivo* chemical environment with reasonable amount of cost, time and labor.^19^ To preserve the original concentration of nutrients and factors in CVF for culture-dependent studies, we collected undiluted CVF in small amounts using Softdiscs. We observed the growth of *L. iners* in microdroplets containing pooled CVF collected from donors with a high prevalence of *L. crispatus*. This work demonstrates the feasibility of collecting and utilizing CVF for high-throughput cultivation of vaginal bacteria. Such a framework opens numerous future directions for culture-based investigation of phenotypes, mechanisms, and inter-species interactions that occur *in vivo*. Our work also establishes an example of incorporating liquids from the native environment, particularly small-volume host fluids, into microdroplet-based microbial cultivation, which can be adapted and extended for the study of a wide variety of complex microbiomes.

## Results

This work describes a new approach for cultivating vaginal bacteria in microdroplets using cervicovaginal fluids (CVF) as the cultivation medium. In particular, we demonstrated the feasibility and potential of this approach by growing *L. iners*, a non-conventional and hard-to-cultivate vaginal lactobacillus, in CVF and comparing its growth to that in laboratory media. As shown in Figure 1, we started by collecting vaginal samples from women of reproductive age using vaginal swabs and Softdisc discs, the latter of which contains both bacteria and CVF. The bacterial composition of each sample was characterized through sequencing of the 16S rRNA gene. Samples collected using Softdisc discs were processed to retrieve cell-free liquids, which were then classified and pooled selectively based on the bacterial composition of the corresponding samples. These liquids were then used as the culture medium to grow vaginal bacteria in nanoliter microdroplets. We observed that *L. iners* ATCC 55195 was able to grow in CVF pooled from samples dominated by *L. crispatus*, which showed significant differences in growth rate and morphology compared to that in a widely used laboratory medium.

**Figure 1:**
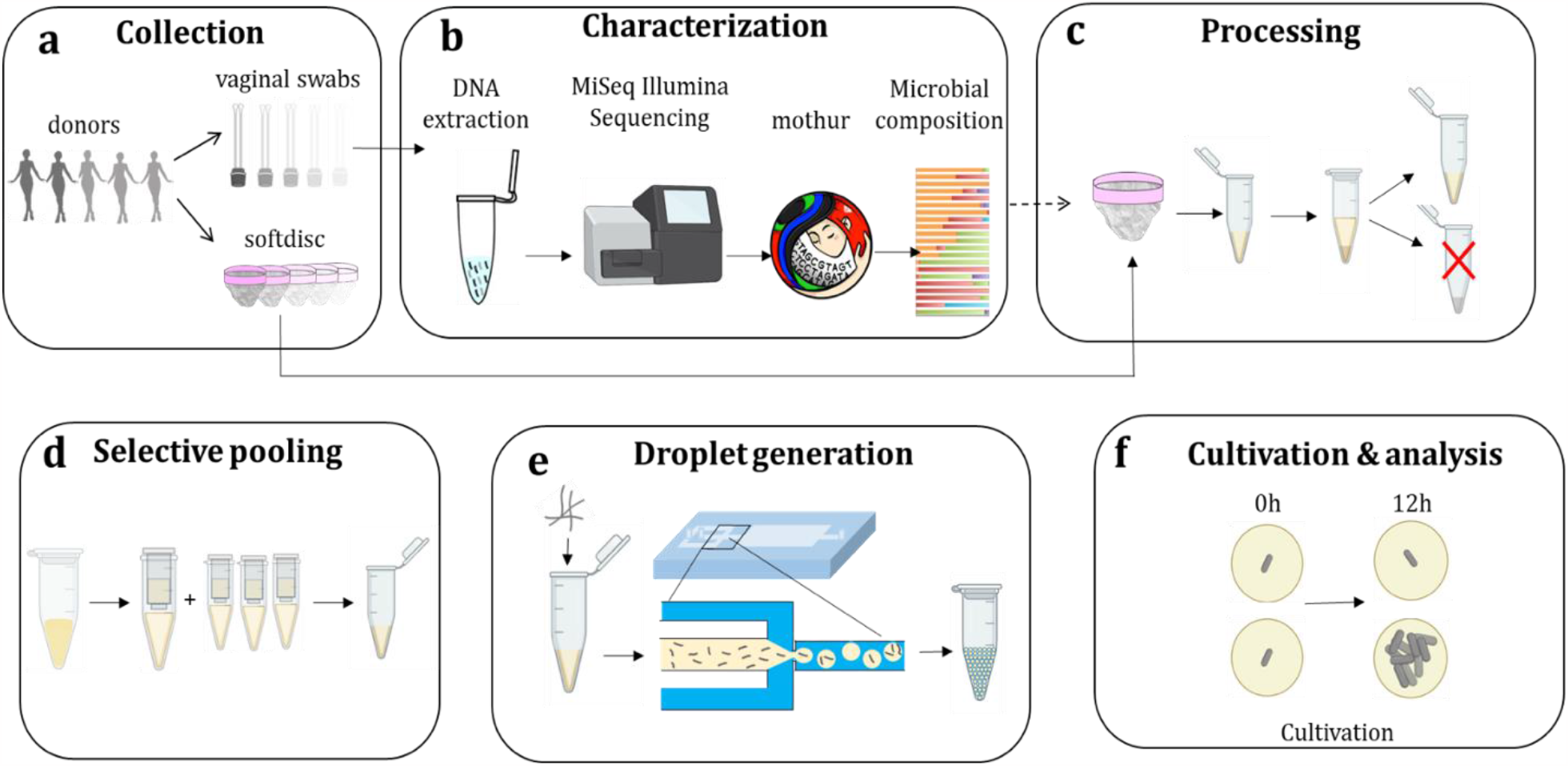
Collection, characterization, processing and pooling of vaginal samples for bacterial cultivation in microdroplets. **a**. Collection of vaginal samples using swabs and Softdisc discs. **b**. Characterization of vaginal bacterial community composition through 16S rRNA genes sequencing. **c**. Processing of CVF-containing samples. Select CVF samples were transferred from Softdisc discs to microcentrifuge tubes. Centrifugation separated the aqueous phase and solid phase. **d**. Aqueous CVF samples were sterilized using 0.22μm filters and pooled selectively. **e**. Vaginal bacteria were spiked into pooled CVF and injected into a microfluidic device with a flow-focusing geometry to generate microdroplets. **f**. Cultivation and analysis of vaginal bacteria in CVF in microdroplets. Solid arrows represent flows of operating steps in a process or flows of materials. The dashed arrow represents the flow of information.

### Subject enrollment, collection and characterization of vaginal samples

A total of seventeen women of reproductive age donated samples in this study. From these donors, a total of thirty-one samples, each consisting of a vaginal swab and a Softdisc disc, were self-collected over a time period of ten months. The number of samples donated by each donor averaged two and ranged from one to seven (Figure S1).

DNA from microbial contents in all thirty-one vaginal swabs and four select Softdisc discs were submitted for 16S rRNA gene amplicon sequencing. The composition of each bacterial community from the swabs are shown in Figure 2 (see Figure S2 for the number of sequencing reads). *Lactobacillus crispatus* was the most frequently detected species, accounting for 66.70 ± 35.24% of each community across all thirty-one vaginal swabs, which is consistent with results from previous surveys of the human vaginal microbiome^20,21^. Twenty-six of these swabs had an abundance ≥ 80% of *L. crispatus. L. gasseri* was the most abundant species in one swab, averaging 1.95 ± 9.11% across all swabs. *L. iners* had an abundance ≥ 80% in two swabs, averaging 11.03 ± 24.34% across all swabs. Low levels of *L. jensenii* were detected for all swabs, averaging 3.57 + 5.88%. Other lactobacilli were detected at low levels across all swabs, averaging 0.35 ± 0.53%. Non-*Lactobacillus* species had an abundance ≥ 80% in three swabs, averaging 16.40 ± 26.66% for all swabs.

**Figure 2:**
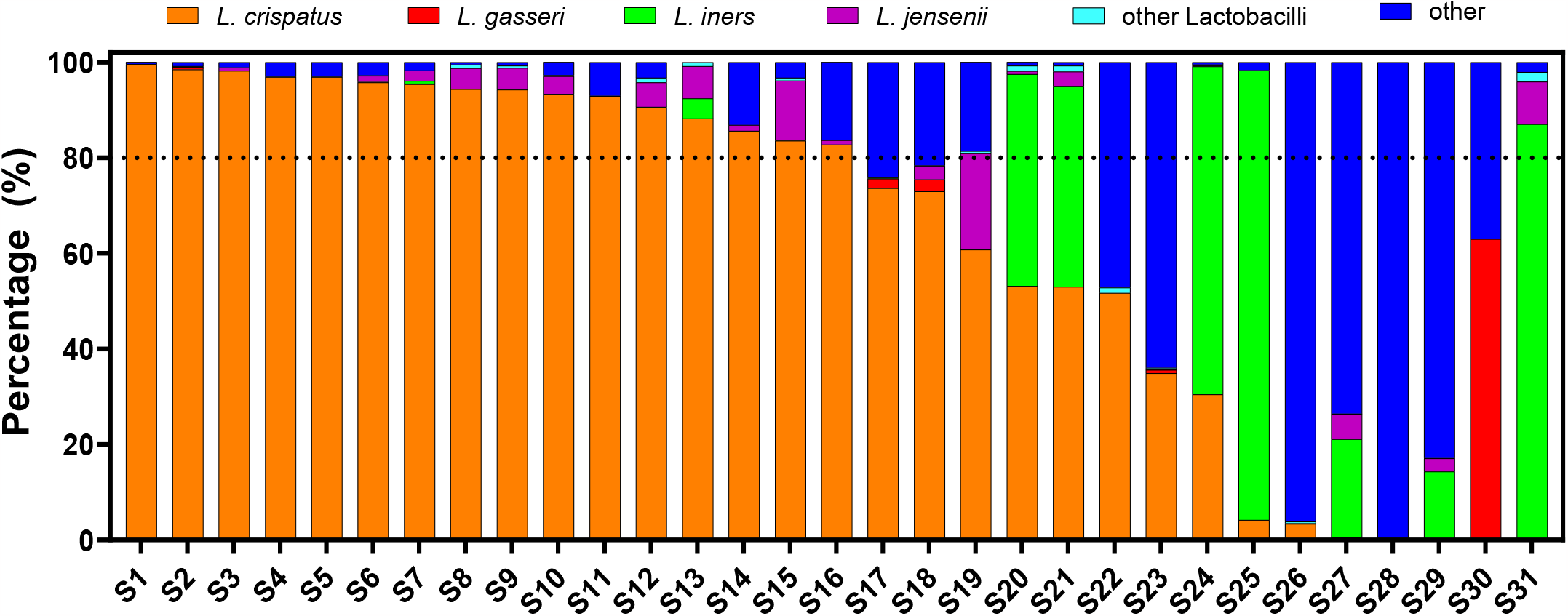
Relative abundance of distinct members of bacterial communities from 31 vaginal swabs.

Sequencing results from four select Softdisc discs were compared to those from corresponding vaginal swabs. As shown in Figure 3 regarding the relative abundance of distinct members (see Figure S2 for the number of sequencing reads), the bacterial communities in the vaginal swab and in the CVF collected with Softdisc at the same time from the same woman were very similar. This result is consistent with previous findings that the microbial community composition in a woman’s vagina and cervix are in high agreement with each other.^22^ Therefore, we decided to rely on results from the vaginal swabs for the microbial composition and conserve CVF as much as possible for subsequent processing in preparation for cultivation of vaginal bacteria.

**Figure 3:**
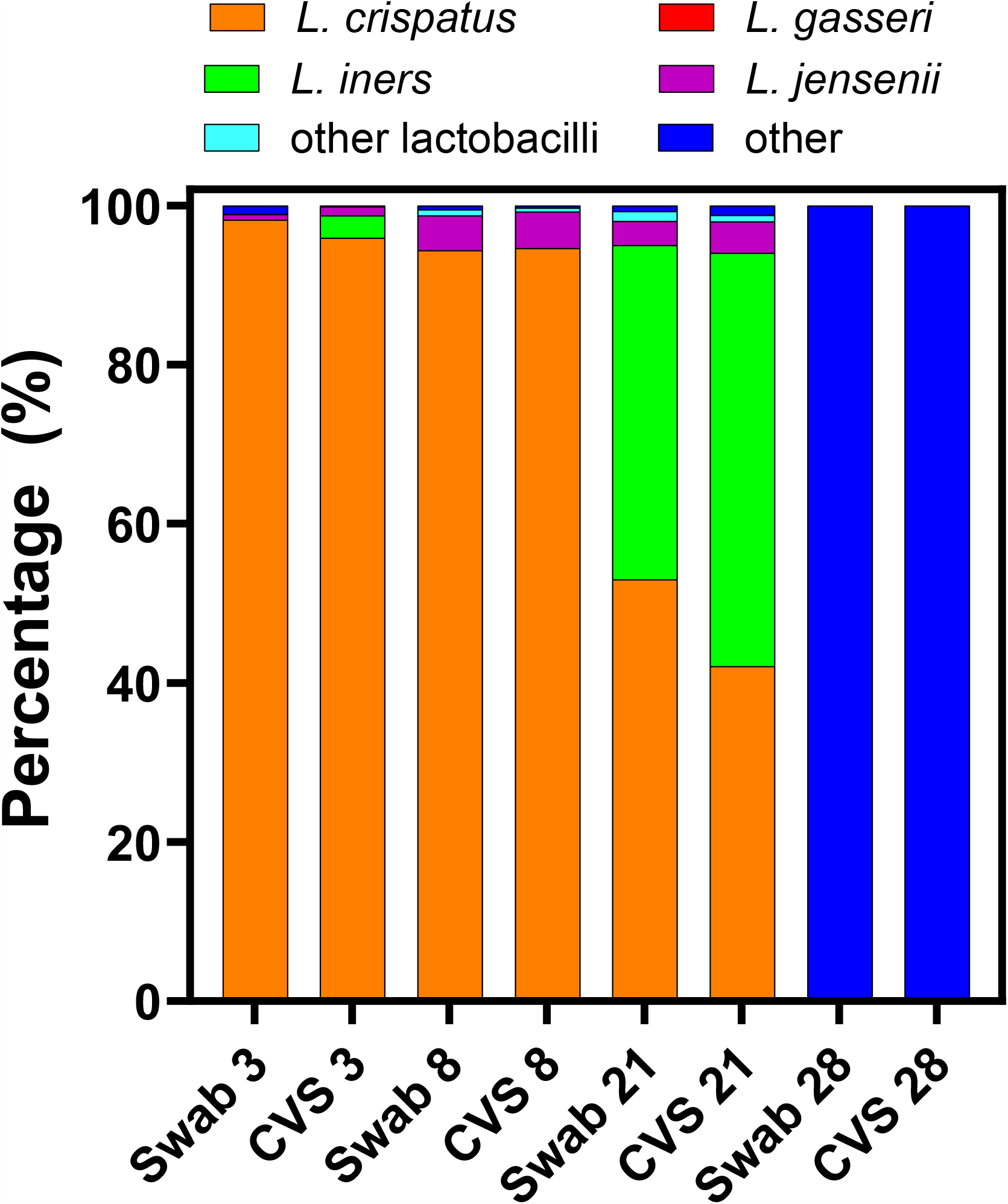
Relative abundance of distinct members of bacterial communities in four sets of vaginal swabs and Softdisc discs (i.e. CVS: cervicovaginal secretion).

We aimed to test the utility of CVF in cultivating vaginal bacteria in microdroplets. To increase the reproducibility of our experiments, we decided to pool CVFs to create an inventory sufficient for multiple experiments. In addition, we decided to categorize CVFs based on the microbial community composition before pooling. As described above (Figure 2), *L. crispatus* was the most abundant species in a majority of the samples.

We thus decided to pool the CVFs for which the relative abundance of *L. crispatus* in the corresponding bacterial community was over a chosen threshold of 80%. A total of sixteen CVF samples met this criterion and each was sterile filtered using a 0.22 μm filter. The volume and pH of these sixteen filtered *L. crispatus* - dominated CVF (LC-CVF) samples are shown in Figure 4. The volume ranged from 5.5 μL to 872 μL, with an average and median of 218.7 ± 233.8 μL and 183 μL, respectively. The pH was found to be in a narrow range of 3.5-4.5, with most samples at ∼4 and an average of 4.0 ± 0.2. To avoid bias from large variations in the volume of these CVF samples, we decided to pool together the same amount from each CVF sample and included thirteen CVF samples each of which had a volume of over 40 μL. Ultimately, 40 μL from each of these thirteen CVF samples were pooled and 520 μL of pooled LC-CVF was created. Aliquots of this pooled LC-CVF were prepared and stored at -80°C.

**Figure 4:**
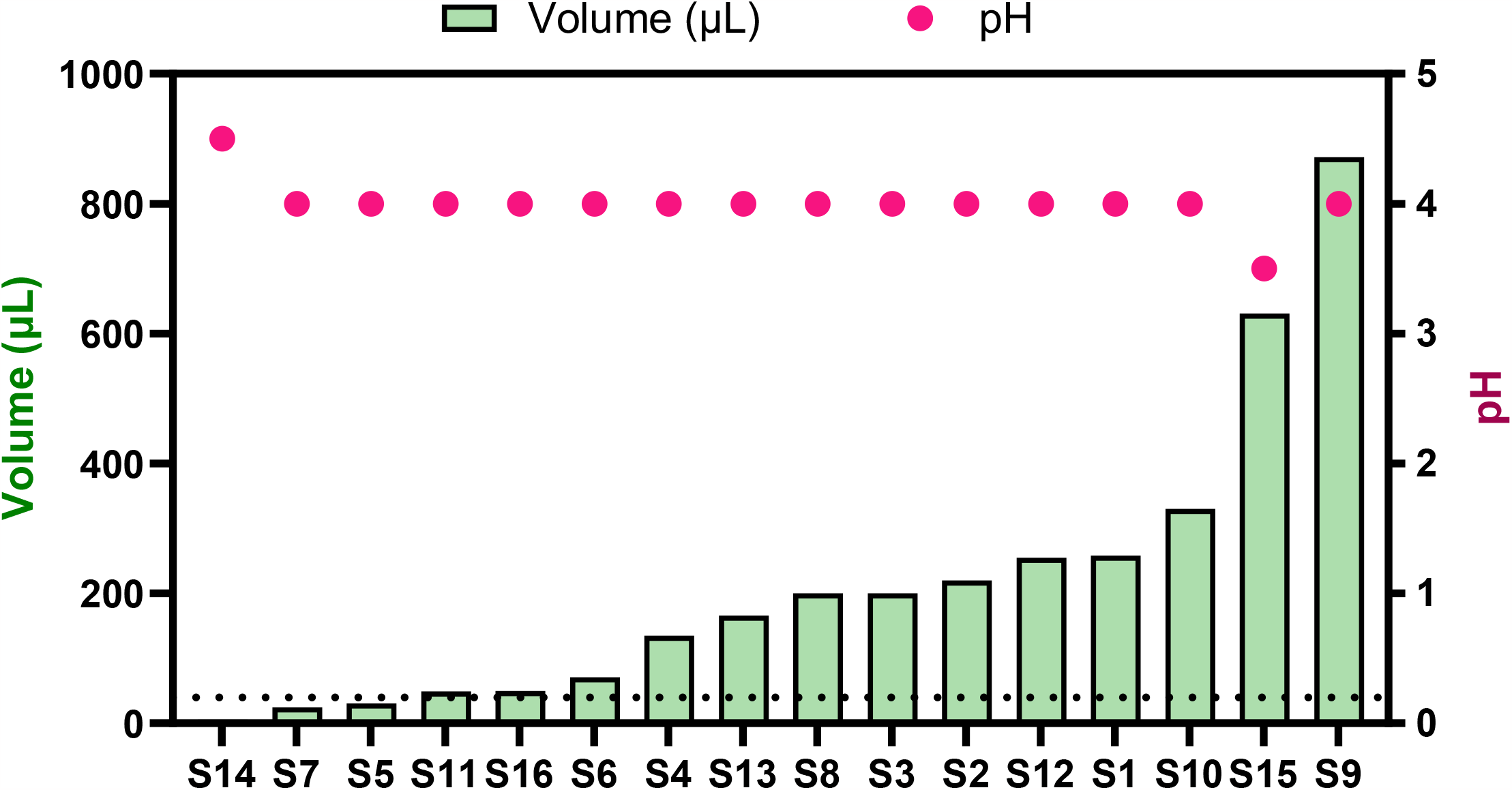
Volume and pH of sixteen sterile-filtered CVF samples with a high prevalence of *L. crispatus*.

### High-throughput growth measurement in droplets with microscopy and image analysis

Droplet analysis must be high-throughput to match the high-throughput of droplet generation and cultivation. To measure growth across hundreds of droplets, we analyzed microscopy images. Using confocal microscopy without fluorescence to quantify cellular growth is possible but requires advanced image analysis techniques (https://www.nature.com/articles/nmicrobiol201677, https://onlinelibrary.wiley.com/doi/full/10.1111/mmi.13264, https://www.nature.com/articles/s42003-019-0480-9). Especially for bacterial growth in droplets, it is difficult to obtain images decipherable by typical image processing algorithms for meaningful high-throughput quantification. However, many vaginal bacteria grow in biofilms. These biofilms are much easier to identify with imaging processing due to their larger sizes and distinct boundaries. We utilized this advantage to quantify the extent of cellular growth in the droplets by measuring the fraction of the droplet area occupied by the biofilm, referred as the biofilm area occupancy. One major disadvantage with this approach is that these biofilms are three-dimensional structures, but microscopy only incorporates the two-dimensional projection. To test if biofilm area occupancy is a good proxy for growth, we correlated biofilm area occupancy with cellular abundance determined by qPCR. *L. iners* was grown in NYCIII media in microdroplets over 28 hours with an initial lambda of 10, with images taken and droplets sampled over time for cell quantification by image analysis and qPCR, respectively. The results show high correlation (R2 = 0.993) for growth up to 14 hours and good correlation (R2 = 0.895) for growth up to 28 hours (Figure S5), showing the utility of biofilm area occupancy metrics from image analysis for highly-parallel growth measurement in droplets.

Further technical improvement would be to incorporate automated imaging to take higher resolution, higher magnification images of single droplets. Recent image analysis packages to quantify high resolution images would allow for the more generalizable study of non-biofilm forming bacterial cultures in droplets.

### Assessment of growth of *L. iners* in different cervicovaginal fluid conditions

Studying the physiological differences between the four major vaginal *Lactobacillus* species is necessary for understanding their exclusion or co-existence. To demonstrate our methodology’s ability to study physiology in ecologically relevant environments, we grew *L. iners* in CVF with different pH. Due to the fastidious nature of *L. iners*, studying its physiology and ecology is particularly difficult among the four *Lactobacillus* species. pH was chosen as an environmental condition to study because it is a critical abiotic factor in vaginal health; CVF is typically slightly acidic but is prone to variations in pH ranging from 4.0 to above 5.5 and may be a critical factor for outcomes regarding interactions of the microbiome with the host. Droplets were generated at 100 μm diameter (105 μm ± 2.8 μm standard deviation) with a lambda of 10 cells/droplet with various media types, such as CVF and the typical laboratory medium used for *L. iners* (which is typically at pH 7.0), each with two pH, 4 and 7. Generated droplets were cultivated anaerobically, and growth was determined using microscopy and image analysis after 24 hours. *L. iners* grew quite well in NYC III pH 7, given it use as the typical laboratory medium and its composition. With only pH adjustments and no further additions, *L. iners* grew moderately well in LC-CVF at pH 7, and very minimally in LC-CVF at pH 4 (Figure 5). Cultivation at pH 4 in NYCIII was also performed, but due to lack of biofilm formation, the image analysis was unable to assess accurate degrees of growth. There was a significant degree of variation across droplets in the same conditions, necessitating the number of droplets needed for proper experimentation. In addition, multiple droplet cultivation experiments were performed with *L. iners* in CVF with pH 4 and 7. These trials demonstrated the same conclusion of higher growth in NYCIII pH 7 than in CVF pH 7, but with wide variability from trial to trial (Figure S7). Some of this variability may be due to the fastidious and sensitive nature of *L. iners*.

**Figure 5:**
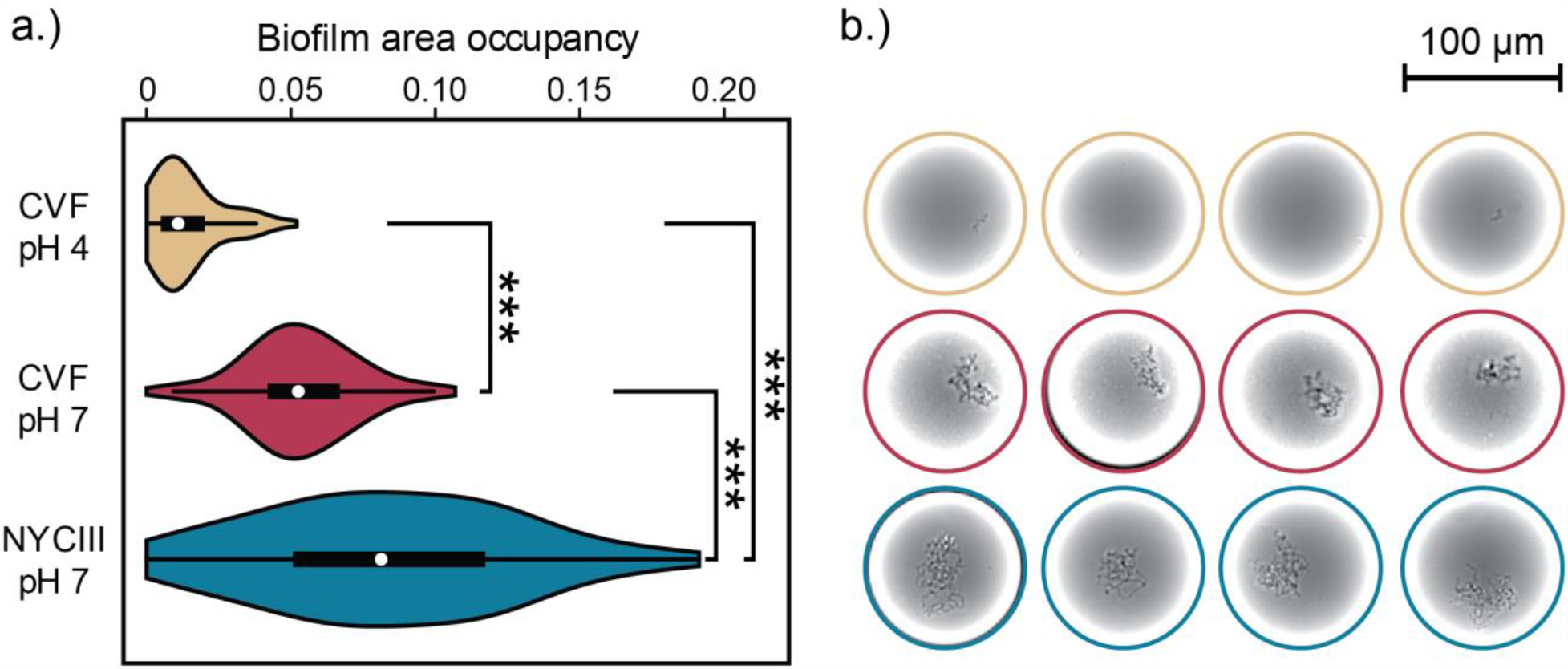
Cultivation of *L. iners* in different growth media in microdroplets. a) Violin plot showing biofilm area occupancy. The three media are: (1) LC-CVF pH 4: cervicovaginal fluid with the native pH of ∼4, (2) LC-CVF pH 7: cervicovaginal fluid with an adjusted pH of ∼7, and (3) NYCIII pH 7: rich NYCIII medium adjusted to pH 7.0. Droplets were incubated for 24 hours, imaged, and analyzed by custom MATLAB scripts to determine what portion of the droplet area was occupied by the *L. iners* biofilm. (b) Representative droplets with biofilms. Top row shows *L. iners* in LC-CVF at pH 4. Middle row shows *L. iners* grown in LC-CVF at pH 7. Bottom row shows *L. iners* grown in NYC III at pH 7. The average initial number of cells in each droplet was 10.

## Discussion and conclusion

In this work, we have adapted a previously developed microdroplet-based high-throughput cultivation platform for the investigation of vaginal bacteria using CVF as cultivation medium.

We observed the growth of *L. iners* in microdroplets containing pooled CVF collected from donors with a high prevalence of *L. crispatus*. This is significant as in previous literature *L. iners* was cultured only in rich laboratory medium. It is also interesting that we observed stronger growth of *L. iners* at neutral pH, which suggests that vaginal pH influences this key species’ growth *in vivo*. On the other hand, it is worth noting that CVF associated with high abundance of *L. crispatus* did not support the growth of *L. crispatus* in our experiment, somewhat surprisingly. This may be due to depleted nutrients or growth factors.

While it is difficult to extend the specific findings about a single bacterial isolate in CVF collected in a single study to the understanding of the vaginal microbiome, the capability demonstrated by this proof-of-concept work is very relevant for the vaginal microbiome, for which the host environment modulates the individual members’ physiological responses and their interactions with one another. Because the analysis taken here is microscopy-enabled, it may also be possible to analyze the interactions of strains in different CVF environments, as long as the strains are uniquely fluorescently-labelled. In terms of medical translatability, this may be used to study how certain key microbial taxa are successful in inhibiting BV-associated anaerobic bacteria in some women but unsuccessful in others. Specifically, fluorescently-labelled *Lactobacillus* strains and BV-associated strains can be co-cultivated together in droplets in CVF samples collected from individual women, assessing if inhibition is observed or not, and performing metabolomic analysis on samples to elucidate which factors may contribute to the different interaction outcomes.

## Material and methods

### Recruiting donors

The University of Michigan Institutional Review Board-Health Science and Behavioral Sciences approved our study, which included recruiting materials like pre-screening questionnaires, flyers, oral scripts, and email messages. All methods relating to recruitment and participation of donors, and use, handling, and storage of vaginal samples were documented in study ID: HUM00111306, titled “Pilot study for microdroplet co-cultivation and analysis of bacterial interactions in the vaginal microbiome.”

Prospective donors were pre-screened through a questionnaire. Healthy female college students, faculty or staff who were i) between the ages of 21 to 45 years old, and ii) were regularly menstruating with a cycle between 25 to 35 days were eligible to participate in the study if they did not meet any of the exclusion conditions that might change the composition of the vaginal microbiome. These exclusion conditions include: 1) recently gave birth, had a miscarriage, or an abortion; 2) using an intrauterine device; 3) is currently pregnant; 4) has had Toxic Shock Syndrome; 5) has had recent gynecological surgery (i.e. Leep, Laser, or cone biopsy) and as a result, has been advised to avoid using internal sanitary tampons or Softdiscs; 6) currently menstruating or using contraceptives interfering with the vaginal mucosa; 7) has taken antibiotics in the past 30 days; 8) has had sexual activity or vaginal discharge within the past 48 hours; and 9) has douched, used vaginal medication, feminine sprays, genital wipes, or contraceptive spermicides. A written informed consent was obtained from each participating woman.

### Collecting vaginal samples

Vaginal samples were self-collected in the women’s restroom at the North Campus Research Complex (NCRC) at the University of Michigan – Ann Arbor. It has been reported that self-collected vaginal swabs have microbial compositions very comparable to those from samples collected by a physician.^29^ In this study, bacterial samples were collected using sterile rayon dual-headed swabs from the Fisherfinest^®^ Dry Transport Swab (Hampton, NH, USA). CVF was collected using an Instead Softdisc™ disc (Venice, CA, USA). Donors were provided with a brown paper bag containing a sterile rayon-tipped dual-headed vaginal swab, an Instead Softdisc disc, a 50 ml Falcon tube, and related instructions. The vaginal swab collection tube and the 50 ml Falcon tube had the same identification number on them for matching the two samples collected from the sample donor. Donors were instructed to wash their hands, enter a bathroom stall, and self-collect samples. Per the instructions, donors unpeeled the swab package, removed the swab without touching the soft tip or laying it down; if the swab came in contact with a substance other than the vagina, donors were instructed to use a new swab. The swab was removed and inserted into its transport tube and placed back into the brown paper bag. To collect CVF, donors washed their hands, re-entered the bathroom stall, and inserted the Softdisc disc following instructions based on this website: http://softdisc.com/how-it-works/. Donors washed their hands after insertion and either set a timer for 15 minutes or returned with the Softdisc disc in less than 12 hours. When the time was up, the Softdisc disc was removed and placed into a 50 ml Falcon tube. The tube was returned to the brown paper bag, which was then placed on dry ice in a Styrofoam cooler located in the same women’s restroom for temporary storage. For long-term storage, all donated vaginal samples were placed in a -80°C freezer until they were processed. All samples were de-identified. Donors were allowed to donate samples no more than once a month and were compensated for each donated sample. Permission to collect vaginal samples in the women’s restroom at NCRC was granted by NCRC, the College of Engineering, the Department of Occupation Safety Environmental Health, and the IRB referred above.

### Processing Softdisc discs

Frozen Softdisc discs in 50ml Falcon conical tubes were thawed on ice at room temperature for 15 minutes. The Falcon tubes were centrifuged at 6,000 rpm for two minutes at 4°C in an Eppendorf Centrifuge 5810 R from Fisher Scientific (Waltham, MA, US) to separate the biological samples from the Softdisc discs. The tubes were then placed back on ice with the bottom deep into the ice to maintain the integrity of the samples inside. These tubes with CVF samples, a Wiretrol® I pipette from Drummond Scientific (Broomall, PA, US), Kimtech^®^ Wipes, sterilized scissors, and sterile 1.5 ml microcentrifuge tubes were placed inside of a biological safety cabinet. From each Falcon tube, the Softdisc disc was removed. The CVF and cervicovaginal mucus (CVM) that remained in the Falcon tube were transferred to a sterile 1.5 ml microcentrifuge tube through the usage of a Wiretrol® I pipette. The Softdisc disc and Falcon tube were disposed of. The 1.5 ml microcentrifuge tube was weighted on a digital gram scale and centrifuged in an accuSpin Micro 17 microcentrifuge from Fisher Scientific at 6,000 rpm for two minutes. The CVM was pelleted at the bottom of the microcentrifuge tube. The CVF was transferred to another sterile 1.5 ml microcentrifuge tube through a pipette. The pH was measured using pH indicator strips by HYDRION^®^ Lab 325 Tape pH Paper Strip Range 3.0-5.5. pH and volume (μL) were recorded.

### Processing vaginal swabs and 16S rRNA gene sequencing

Vaginal swabs were placed in a cooler with dry ice and transported to the Medical Research Science Building I at the University of Michigan – Ann Arbor. There the vaginal swabs were placed into a biological safety cabinet along with a Glass Bead Sterilizer, three pet nail trimmers, two sterile glass Petri dishes to prop the trimmers, and a 96-well PowerMag Glass Bead plate (Qiagen, Hilden, Germany). The Glass Bead Sterilizer was turned on. When it was ready, the trimmers were placed in the hot glass beads and sterilized. A soft-tip from each vaginal swab was clipped and dispensed into a separate well in the 96-well plate. After all swabs were processed, the plate was sealed, packaged, and submitted for 16S rRNA gene amplicon sequencing according to instructions at the University of Michigan Microbial Community Analysis Core. The core carried out gDNA extraction, polymerase chain reaction (PCR) using a Dual indexing sequencing strategy, and 16S rRNA gene sequencing using Illumina’s MiSeq platform. DNA extraction was carried out with the Eppendorf EpMotion liquid handling system and the Qiagen MagAttract PowerMicrobiome kit (previously MoBio PowerMag Microbiome). The V4 region of the 16S rRNA gene was amplified and sequenced. The Dual-index sequencing strategy was developed by Kozich et al..^30^ The PCR protocol was as follows: Stage 1, 1 cycle of 95°C (2 min); Stage 2, 30 cycles of 95°C (20 s), 55°C (15 s) and 72°C (5 s); and Stage 3, 1 cycle of 72°C (10 min); Stage 4, 1 cycle of 4°C (forever). Each PCR mix (20 μL) was composed of 2 μL 10x AccuPrime PCR Buffer II, 0.15 μL AccuPrime HiFi Polymerase, 1 μL DNA, 5 μL primer set (4 μM) and 11.85 μL water.

### Analyzing 16S rRNA gene sequencing data

The V4 region of 16S rRNA gene sequences was identified as described by Chakravorty et al.^31^ and Yarza et al.^35^ Mothur, an open-source software for bioinformatic analysis of microbial communities,^32^ was used to process the 16S rRNA gene sequences. We input sequences from the SILVA database for *Lactobacillus* species and established a consensus sequence of the V4 region for each vaginal *Lactobacillus* species. Sequences were aligned based on methods described on the mothur blog entitled “Customize your reference alignment for your favorite region” (https://mothur.org/blog/2016/Customization-for-your-region/). The resulting operational taxonomic units (OTUs) were categorized into six groups: *L. jensenii, L. iners, L. gasseri, L. crispatus*, other *Lactobacillus*, and others that are within the genus of *Aerococcus, Atopobium, Bifidobacteriaceae, Clostridia, Corynebacteriume, Dialister, Finegoldia, Megasphaera, Mobiluncus, Peptoniphilus, Prevotella*, and *Veillonella*. These genera were detected in the vagina in previous studies.^33,34^

### Bacterial strain and cultivation

*L. iners* ATCC 55195 was originally isolated from the human vagina.^28^ For starting a culture, cells from a cryostock were streaked onto Tryptic Soy Agar (TSA) with 5% sheep blood (Thermo Scientific, R01198) and incubated for two days in an anaerobic chamber (with 2-3% hydrogen, 10% carbon dioxide, balance nitrogen) at 37°C. Single colonies from the plate were inoculated in 1 mL volumes of NYCIII medium (referred from ATCC medium 1685, per 1 L: HEPES, 4.0 g; proteose peptone No. 3, 15.0 g; glucose, 5.0 g; yeast extract, 3.75 g; heat inactivated horse serum, 100.0 mL; adjusted to pH 7.3) and incubated anaerobically at 37°C for 12-16 hours to grow cells within early log phase. 1 mL of turbid culture was washed three times in PBS (centrifugation at 4000x for 5 minutes, removal of supernatant, and resuspension in PBS). Cells were diluted and counted with a disposable haemocytomer (C-Chip: SKC, Inc. DHCN015). Cells were inoculated in the appropriate media to achieve a λ value (the average number of cells per microdroplet) of 10 (around a 10-fold dilution) for 100 μm diameter microdroplets.

For microdroplet generation, two media were used: NYCIII growth medium and *L. crispatus*-dominated CVF with two conditions for each, pH 4 and 7. Without any modification, CVF measured around pH 4 using pH paper. To adjust CVF, due to the small volumes (typically 20-50 uL per trial), we added increments of 0.1 uL 5 M NaOH, mixed with pipetting, and checked the pH with pH paper until pH 7 was reached.

### Microfluidic device fabrication

Microfluidic devices used to generate microdroplets were made with polydimethylsiloxane (PDMS). Devices were made by mixing uncured PDMS base elastomer and curing agent (10:1 mass ratio of elastomer to curing agent) and pouring the mix onto SU-8 molds with the microfluidic device features (Figure S4). Air bubbles were removed in a vacuum chamber and the PDMS was cured overnight at 65°C. The cured PDMS layer was peeled off the SU-8 mold, and devices were cut to size. For inlets and outlets to the device, holes were punched with a biopsy punch (1.0 mm inner diameter). PDMS devices were bonded on glass microscope slides via plasma-activated bonding using a corona discharge wand and silanized with (tridecafluoro-1,1,2,2,-tetrahydrooctyl)-1-trichlorosilane using a desiccator.

### Microdroplet generation and cultivation in microdroplets

Microdroplet generation requires two co-flow phases: an oil phase with emulsion-stabilizing fluorosurfactant and an aqueous phase suspension with cells and media. The oil phase was Novec HFE-7500 fluorinated oil (3M) with 2% PEG-PFPE amphiphilic block copolymer surfactant (Ran Biotechnologies, 008-FluoroSurfactant). The aqueous cell suspension was the cell suspension in growth media (NYCIII or CVF). The aqueous and the oil phase were loaded into 1 mL and 3 mL syringes, respectively, with 23-gauge Luer Lock syringe needles (BD 305145) attached. Because of the low volume of CVF (20-50 uL per droplet generation event), in order to push the entire volume through the device, the cellular suspension was withdrawn into a syringe filled mostly with air, keeping the suspension in the needle, making sure no air bubbles were intaken. To mix the cellular suspension, the CVF was kept in a 1.5 mL microcentrifuge tube and mixing was done with a micropipette, being careful not to introduce air bubbles. PTFE tubing (0.022” ID, Cole-Parmer) was used to connect the syringe needle to the PDMS device. Kent Scientific GenieTouch syringe pumps were used to infuse the oil phase and cell suspension phase into the device to generate the droplets. For 100 μm diameter droplets, the oil phase flow was set to 30 μL/min and the aqueous flow rate was set to 20 uL/min, allowing the air in the syringe to push the continuous flow of CVF into the device. Droplets were collected in tubes from the outflow of the droplet generation device through PTFE tubing. Mineral oil was placed on top of the droplet emulsion to prevent evaporation of droplets but allow exchange of air. The droplets were then placed in the anaerobic chamber to incubate at 37 C under previously mentioned anaerobic conditions. To visualize growth in droplets, in the anaerobic chamber, 3-5 uL of the droplet emulsion and the remaining volume of oil with surfactant (to 10 uL volume) were pipetted into a C-Chip, and the inlet and outlets were epoxied to prevent evaporation. The C-Chip was removed from the anaerobic chamber and placed under a Nikon Ti-S inverted microscope and photographed with a Retiga R6 camera (Teledyne QImaging).

### Image analysis of biofilm growth

To quantify growth in droplets in a highly-parallel fashion, we used microscopy and image analysis. *L. iners* tends to grow in large clumps which form distinct biofilms. Unlike typical planktonic growth where delineating cell boundaries is difficult, biofilm boundaries can be detected easily with image processing. We developed a MATLAB script utilizing various image process functions to determine what percentage of the droplet area in the image was occupied with biofilm. The MATLAB script is available in the Supplementary Information (make sure I attach it) with the images analyzed for all our droplet cultivation trials. Examples of the image detection are shown in Figure S6. The script detects uses edge detection and dilation to detect biofilms within droplets, and droplets within the acceptable range are selected based on circle detection algorithms and the range set by the user, allowing us to disregard merged droplets.

### Cell lysis and qPCR

Because the size of the biofilm can only be assessed from the 2-D microscopy image, we benchmarked the accuracy for biofilm occupancy (the fraction of the droplet area occupied by the biofilm) as a proxy for growth in the droplet. To do so, we cultivated *L. iners* in NYCIII droplets according the same fashion as previous experiments and quantified growth with image analysis and compared with abundance measured with qPCR. To do so, we started with a λ value of 10 in 100 um diameter microdroplets and incubated them anaerobically, in the same fashion as above. We generated 5 50 uL volumes of droplet emulsion with 20 uL of oil and surfactant with 50 uL of mineral oil on top in 5 separate 0.2 mL PCR tubes. To collect droplets at different timepoints (at 0, 4, 8, 14, 28 hrs), we removed a whole PCR tube from the anaerobic chamber and sampled droplets from this tube and processed them through droplet breaking, cell lysis, and qPCR. For DNA extraction, we processed three replicates for each timepoint, processing 5 uL of droplets for each. For each replicate, we destabilized microdroplets in 1.5 mL microcentrifuge tubes by adding 1 uL of perfluoro-1-octanol and 50 uL of sterile TE buffer (10 mM Tris-HCl, 50 mM EDTA, pH 8) and mixed with vortexing. We performed this for all samples and processed them when all samples were collected with DNA extraction. DNA extraction was a modified procedure from Forney et al. (DNA extraction paper) with only lysozyme in the enzymatic treatment. For each sample, 10 uL of 50 mg/mL lysozyme and 10 uL of TE buffer was added to each replicate and allowed to incubate at 37 C for one hour. We added 5 uL of Proteinase K (need to find conc) and 50 uL Buffer AL, mixed with pulse vortexing for 15 seconds, and incubated for 10 minutes at 56 C. We centrifuged at 10000 g for 1 min and transferred 50 uL of the liquid to a clean 1.5 mL microcentrifuge tube, making sure not to grab oil or perfluorooctanol. To each we added 5 uL of 3 M sodium acetate pH 5.5 and 50 uL of 100% EtOH and mixed by pulse-vortexing for 15 seconds, briefly centrifuging to collect the liquid on the bottom of the tubes. We then placed the samples into silica gel purification EconoSpin columns (Epoch Life, 1920-250) and centrifuged full speed for 1 min. For washing of DNA, we added 450 uL of Buffer AW1 to the column, centrifuged at full speed for 1 min, added 450 uL of Buffer AW2, centrifuged at full speed for 1 min, centrifuged at full speed for 1 min to remove residual liquid, and placed the columns into a new 1.5 mL microcentrifuge tube for elution. For elution, we placed 20 uL of Buffer AE which had been incubating at 60 C into the column, incubated in the column for 5 minutes, and centrifuged the eluate for 1 min. Quantification of DNA was performed with a QuantStudio3 qPCR thermocycler with PowerUp SYBR Mastermix and *L*.*iners* specific 16S PCR primers (Alqumber et al.). Exact PCR mix was as follows, per 20 uL reaction: 10 uL 2X PowerUp SYBR Mastermix, 1 uL of template from DNA extraction eluate, 0.6 uL of each forward and reverse primer, and 7.8 uL of molecular grade water. The thermocycler program was: 50 C for 2 minutes; 95 C for 2 minutes; 70 cycles of: 95 C for 30 seconds, 53 C for 10 seconds, and 72 C for 10 seconds, and 72 C for 10 minutes.

