## Supplementary Data for "Microdroplet-enabled high-throughput cultivation of vaginal bacteria using cervicovaginal fluids"

**Figure S1**: The number of samples from each of seventeen donors. A sample is defined as one vaginal swab and one Softdisc disc. Mean and standard deviation are 1.8 ± 1.0 samples.

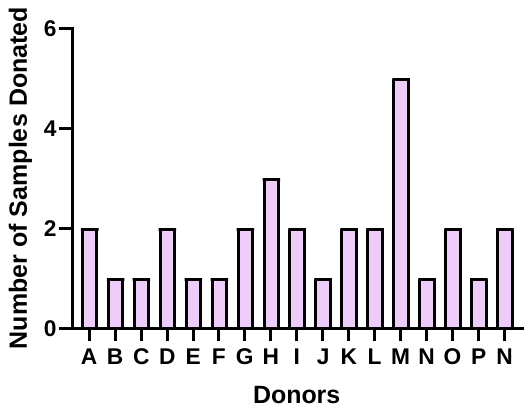

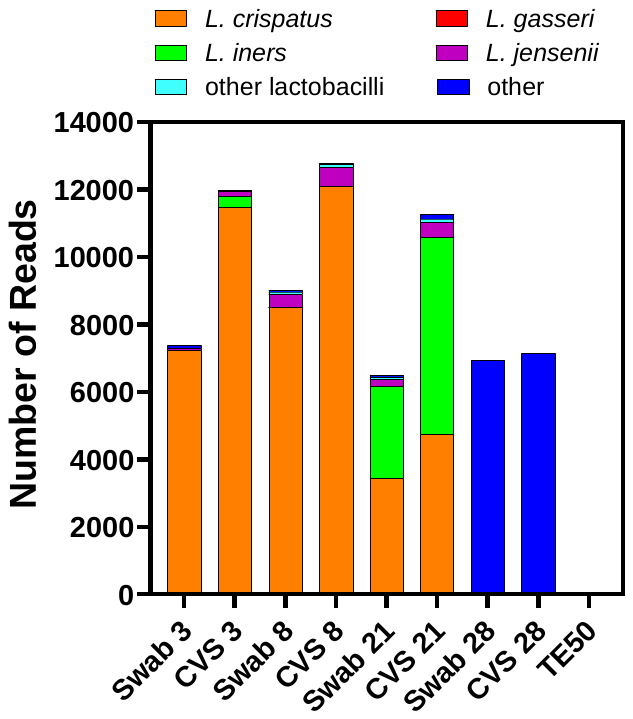

**Figure S3:** The number of sequencing reads for distinct members of bacterial communities in four pairs of vaginal swabs and Softdisc discs. CVS: cervicovaginal secretion.

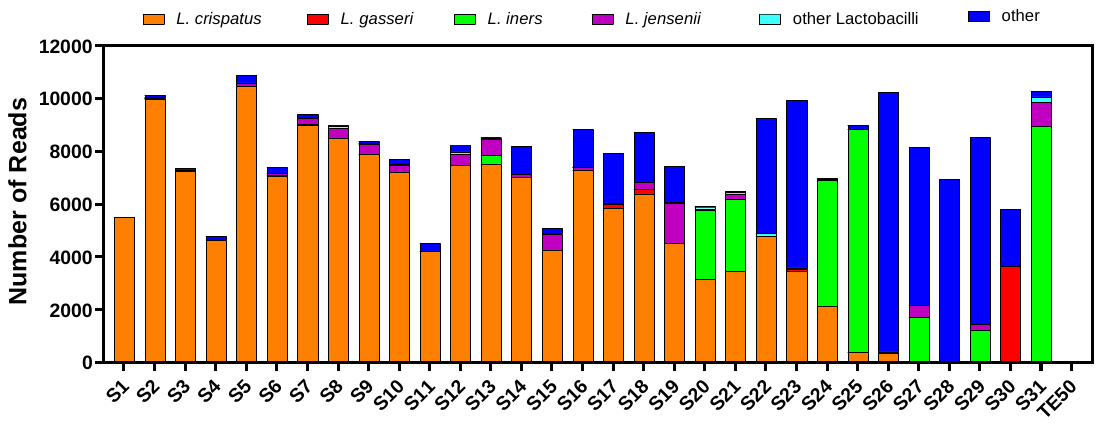

**Figure S2**: The number of sequencing reads for distinct members of bacterial communities from 31 vaginal swabs. The number of reads per vaginal swab ranged from 4,789 to 10,913, and averaged 7,932.

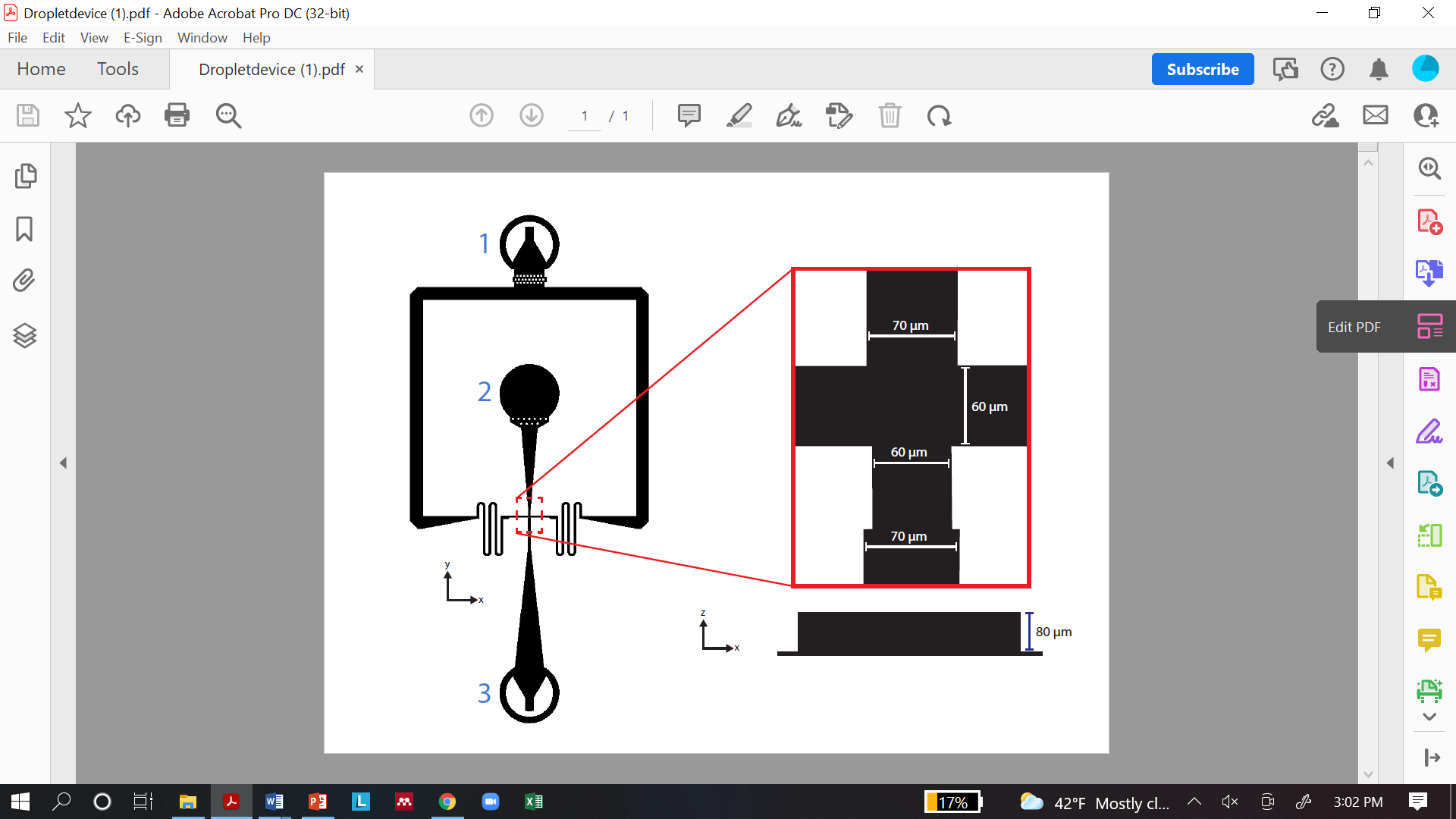

**Figure S4:** Schematic layout of the flow-focusing microfluidic device used for droplet generation**.** The device has an oil inlet (1), cell suspension inlet (2), flow-focusing intersection (boxed in red), and a droplet outlet (3). The intersection (right) has channel widths of 60 or 70 μm and a channel height (blue scale bar) of 80 μm. The device was used to generate droplets with a diameter of ~100 μm.

**Figure S5:** Log fraction of droplet area occupied by cells with respect to Ct value from qPCR over 28 hours. The red dashed line is the line of regression for 0, 4, 8, and 14 hours. The blue dashed line is the line of regression for 0, 4, 8, 14, and 28 hours.

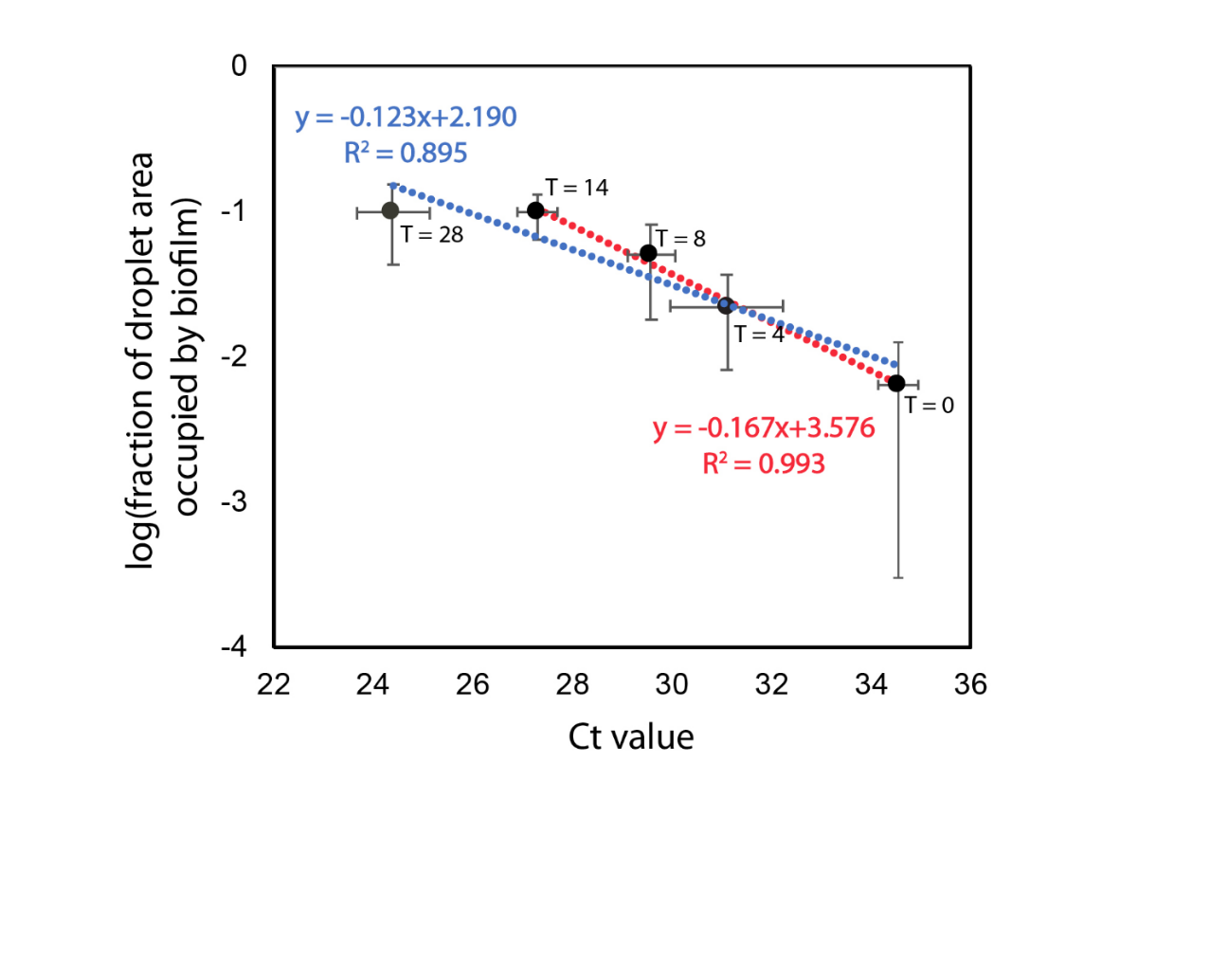

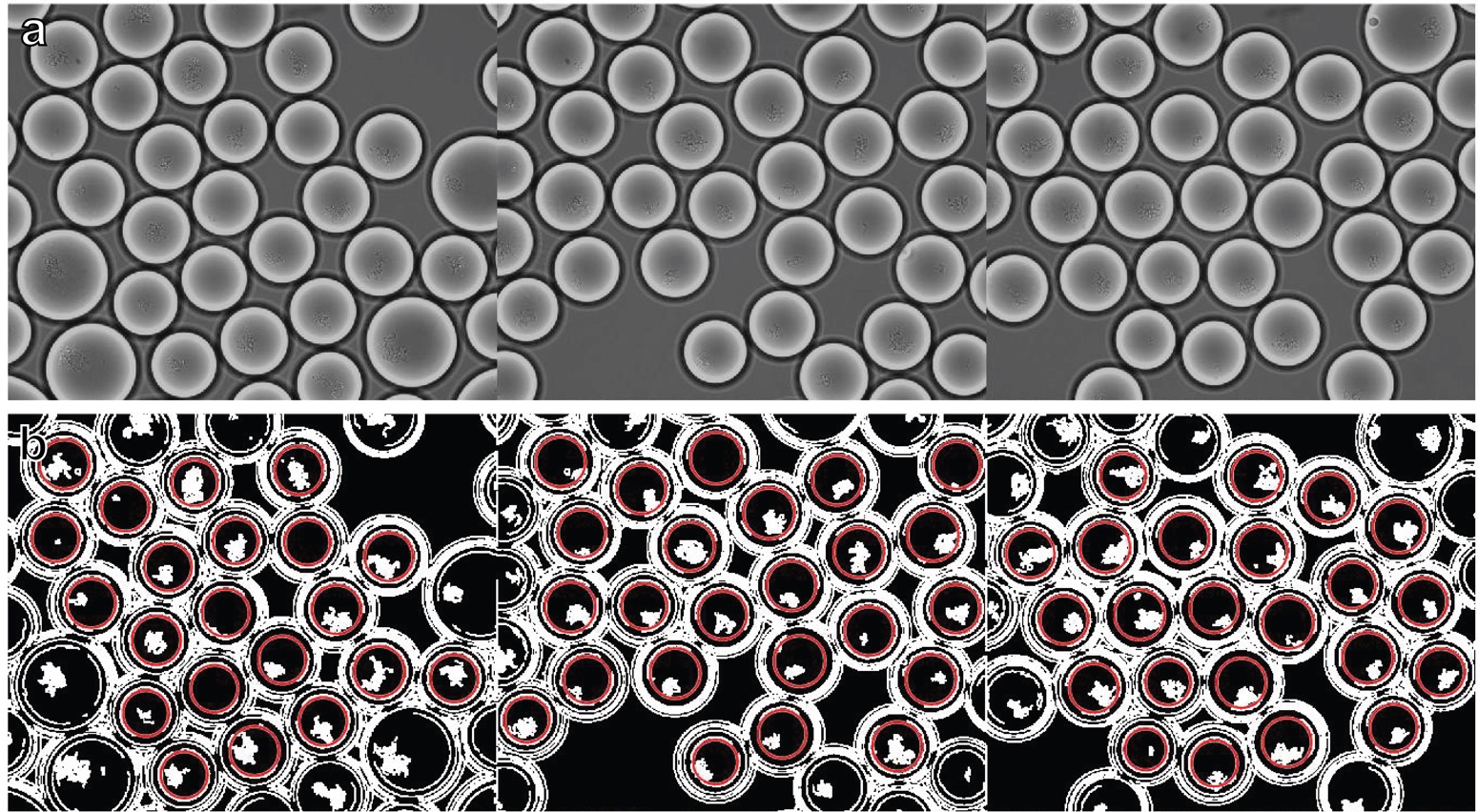

**Figure S6:** Examples of image processing for the quantification of growth in droplets from microscopy images. (Top row) Confocal microscopy images. Average microdroplet diameter is 100 um with merged droplets appearing larger. Biofilms within the droplet boundary are visibly distinct. (Bottom row) Images after processing in MATLAB. Red circles indicate droplet detection, selecting only unmerged droplets. The biofilm boundaries are identified, dilated, and used for quantification.

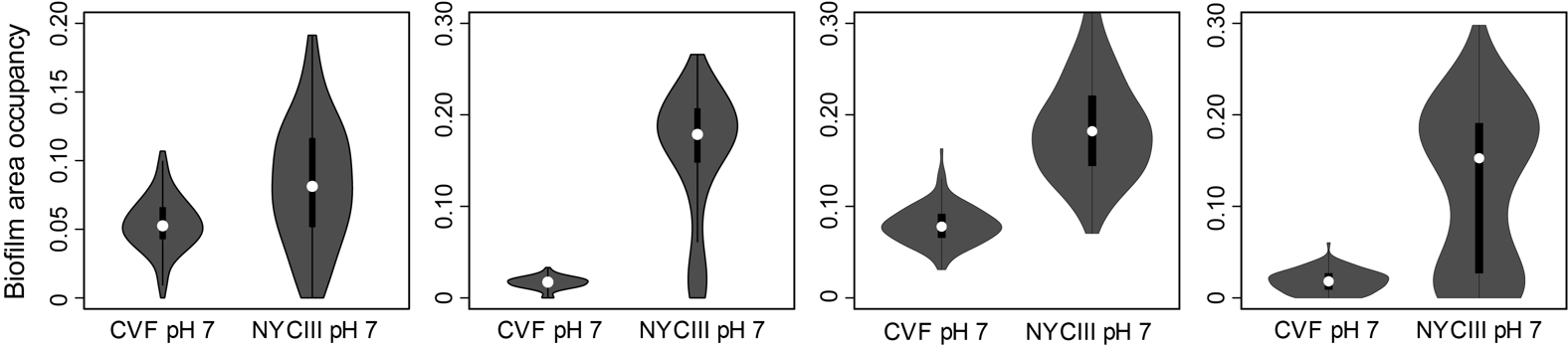

**Figure S7:** Violin plots of *L. iners* growth in droplets (quantified with biofilm area occupancy fraction) after 24 hours from 4 different trials of droplet cultivation of *L. iners* in LC-CVF and NYCIII at pH 7. The NYCIII media and LC-CVF and *L. iners* strain was the same for all trials, but variability in the extent of growth across trials is observed.
